# Characteristic differences in tibial subchondral bone changes in the post-traumatic knee osteoarthritis model

**DOI:** 10.1101/2023.02.27.530166

**Authors:** Kohei Arakawa, Kei Takahata, Yuna Usami, Takanori Kokubun

## Abstract

**Objective:** Meniscus degeneration and subchondral bone changes contribute to the development and progression of knee OA. The purpose of this study was to reveal the relationship between medial meniscus degeneration and characteristics of subchondral bone changes in different models

**Design:** We used the anterior cruciate ligament transection (ACL-T) model, destabilization of the medial meniscus (DMM) model, controlled abnormal tibial translation (CATT) model and a controlled abnormal tibial rotation (CATR) model with different mechanical stresses. We performed histological analysis and micro computed tomography analysis, at 4 and 6 weeks. In addition, we divided the tibial subchondral bone into four compartments and set the regions of interest

**Results:** Meniscus degeneration was observed in all groups, but there was no significant difference. The ACL-T group showed posterior displacement of the contact area, and the DMM and CATR groups showed lateral deviation of the medial meniscus. The region-specific subchondral bone changes in each model showed that changes in mechanical stress due to ACL and meniscus dysfunction, as well as changes in the contact area, affect the bony structure of the subchondral bone differently in each region.

**Conclusions:** Subchondral bone changes in different models of mechanical stress were different in each region. In particular, changes in the contact area and increased compressive stress due to meniscus dysfunction were suggested to promote bone formation. The results of this study indicate that changes in alignment and contact area in the PTOA model may cause region-specific characteristics of the subchondral bone changes.

## INTRODUCTION

Knee Osteoarthritis (OA) is characterized by articular cartilage degeneration, which causes joint dysfunction. Knee OA is a disease of the entire joint, affecting the articular cartilage and surrounding tissues. In particular, the subchondral bone can cushion mechanical stresses with articular cartilage and interacts with articular cartilage.^1–4^ Therefore, articular cartilage and subchondral bone are considered functional units. Post-Traumatic Osteoarthritis (PTOA) is a subtype of OA primarily caused by joint injury and includes anterior cruciate ligament (ACL) tears and meniscus injuries. Since the ACL is responsible for 86% of the anterior tibial translation^5^, anterior-posterior joint instability occurs in patients with ACL tears. In addition, the meniscus is responsible for stabilizing the joint and distributing compressive stresses^6^, resulting in increased compressive stresses in patients with meniscus injuries. Therefore, increased mechanical stress due to ACL and meniscus dysfunction is involved in the onset and progression of PTOA^7, 8^.

The relationship between knee OA progression and subchondral bone changes is thought to be that early-stage knee OA accelerates subchondral bone resorption. In contrast, late-stage knee OA accelerates subchondral bone sclerosis.^9, 10^ In addition, it has been reported that compressive loading on the tibia promotes bone formation^11^ and that the reduction of mechanical stress by hindlimb suspension promotes bone resorption.^12^ Our previous study revealed different characteristics of subchondral bone changes during the early knee OA phase between models that reproduced increased or decreased mechanical stress due to the ACL and meniscus dysfunction.^13^ However, the characteristics of subchondral bone changes are not completely understood, and further studies are needed. The understanding of early changes in the subchondral bone is important for the maintenance of subsequent joint function.

Previous studies have shown that the properties of articular cartilage differ between the area covered by the meniscus and the area uncovered by the meniscus. Especially the area covered by the meniscus is fragile.^14^ In addition, degeneration of the meniscus has also been observed in animal models of knee OA,^15^ and it is thought that the meniscus dysfunction leads to increased mechanical stress on the tibial articular surface. An implication of these cartilage properties is the possibility that the properties of the subchondral bone are also different and may be affected by the increased mechanical stress caused by meniscus dysfunction. In previous studies, we focused on ACL and meniscus dysfunction that affected cartilage and bone differentiation and established some specific models of mechanical stress. Then, we reported the possibility that different mechanical stresses may have different effects on the subchondral bone.^13, 16^ Thus, we hypothesized that subchondral bone changes would depend on increased or decreased mechanical stress and that the mechanical properties would differ in each region of the tibial articular surface. Therefore, the purpose of this study was to reveal the relationship between medial meniscus degeneration and characteristics of subchondral bone changes in different models of mechanical stress that we had established.

## MATERIALS AND METHODS

### Animals and Experimental Design

This study was approved by the Animal Research Committee of Saitama Prefectural University (approval number: 2020-1). The experimental design is illustrated in Figure 1A. In this study, 40 twelve-week-old ICR (Institute for Cancer Research) male mice were procured for the study. The mice were randomized into one of five groups: ACL-transection (ACL-T) group (ACL-T, n = 8), controlled abnormal tibial translation (CATT) group (CATT, n = 8), destabilization of the medial meniscus (DMM) group (DMM, n = 8), controlled abnormal tibial rotation (CATR) group (CATR, n = 8), and sham group (Sham, n=8). All mice were housed in plastic cages under a standard 12 h light/dark cycle. Mice were permitted unrestricted movement within the cage and had free access to food and water.

**Figure 1.**
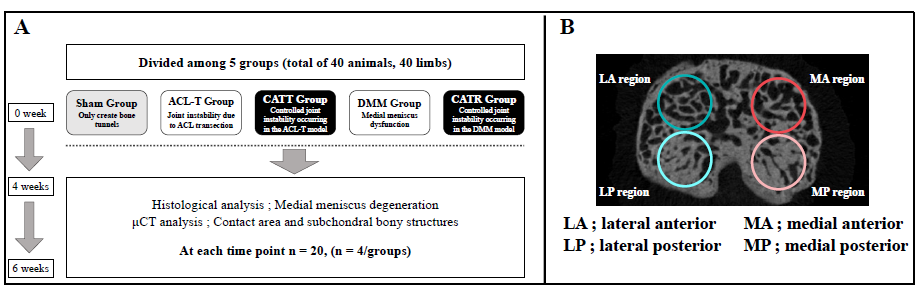
Experimental design and regions of interest (A) Experimental design. We performed histological analysis and μCT analysis at 4 and 6 weeks. These analysis involved the Sham group, ACL-T group, CATT group, DMM group, and CATR group (for each group, n = 4). (B) Subchondral bone regions of interest. We divided the tibial subchondral bone into four regions (MA, MP, LA, and LP regions) and performed μCT analysis.

### Surgical Procedures

The surgical procedures were performed with reference to our previous study.^13, 16^ ACL-T surgery, the medial capsule was exposed, and scissors were used to cut the ACL. In the ACL-T group, shear stress increases with joint instability due to ACL tear. DMM surgery, the medial capsule was exposed, and scissors were used to cut the medial meniscotibial ligament (MMTL). In the DMM group, shear and compressive stress increase due to meniscus dysfunction. CATT and CATR surgery followed the same procedure as ACL-T and DMM surgery. These models created bone tunnels in the distal femur and proximal tibia using a 25-gauge needle and 4-0 nylon threads threaded through them. For the position of the bone tunnels, the CATT model was created to follow the running of the ACL. In contrast, the position of the bone tunnels in the CATR model was created more vertically than in the CATT model. The CATT group reduces the increase in shear stress caused by ACL rupture by suppressing joint instability. On the other hand, the CATR group minimizes the increase in shear stress caused by joint instability, and compressive stress increases due to meniscus dysfunction. To cancel the differences in intervention among the groups, a bone tunnel was also created in Sham groups, and a nylon thread was tied loosely to maintain joint instability. Details of surgical intervention methods are presented in Supplementary Methods (Supplementary Fig.1).

**Supplementary Figure 1.**
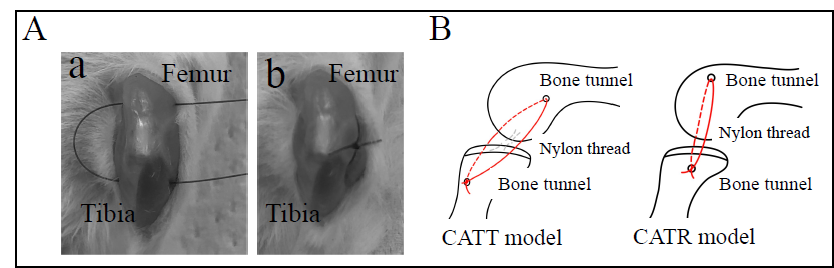
Surgery for the CATT and CATR model. (A) Details of surgical intervention methods. Create bone tunnels in the femur and tibia (a). Nylon threads are threaded through the bone tunnels and tightly tied (b). (B) Schematic diagram of the CATT and CATR models. Nylon suture suppresses anterior-posterior joint instability caused by the ACL-T model and rotational joint instability caused by the DMM model.

### Histological Analysis

Mice were sacrificed at 4 and 6 weeks after surgery. Subsequently, the knee joint was fixed in 4% paraformaldehyde/phosphate-buffered saline for 24 h, followed by decalcification in 10% ethylenediaminetetraacetic acid, dehydration in 70% and 100% ethanol and xylene, and embedding in paraffin blocks. Thin sections (6 μm) were cut in the sagittal plane using a microtome (ROM-360; Yamato Kohki Industrial Co., Ltd., Saitama, Japan), stained with safranin-O/fast green, and subjected to histological evaluation to estimate the degree of medial meniscal degeneration on one slide of each sample. All the samples were taken with a fluorescence microscope BZ-X710(Keyence, Tokyo, Japan), and the meniscus was taken at 20x magnification.

The mouse meniscus histological grading system established by Kwok et al.^15^ was used to evaluate the degeneration in the anterior and posterior horn of the medial meniscus. One independent observer (KT) performed meniscus scoring on a five-point scale (0-4).

### Micro-computed tomography (μCT) analysis of subchondral bone

The knee joints were scanned using a *μ*CT system (Skyscan 1272, BRUKER, MA, USA) with the following parameters: pixel size, 6μm; voltage, 60 kV; current, 165 μA. Subsequently, the reconstructed image was acquired using the NRecon software (BRUKER, MA, USA). We divided the tibial subchondral bone into four compartments and set the regions of interest (medial anterior; MA region, medial posterior; MP region, lateral anterior; LA region, lateral posterior; LP region) (Fig.1B). We then designated volumes of interest on the 40 slides of subchondral bone in each region. Then calculated the bone volume/tissue volume fraction (BV/TV, %) for each region of interest using CTAn software (BRUKER, MA, USA). Subsequently, we compared regions within a model and for comparisons between models per region.

### Statistical Analysis

All analyses were performed using R Studio version 1.2.5019. The normality of the data distribution was assessed using the Shapiro-Wilk test. Subchondral bone data were compared using a one-way analysis of variance (ANOVA), and post hoc Tukey’s test was used. Meniscus scores were compared using the Kruskal-Wallis test and post hoc Steel-Dwass analysis. All significance thresholds were set to 5%. Parametric data are presented as mean ± 95% confidence interval, and nonparametric data are presented as median with interquartile ranges [IQR].

## RESULTS

### Histological Analysis

Medial meniscal degeneration was evaluated 4 and 6 weeks after surgery (Fig.2). We presented the histological image of the meniscus in figure 2. At 4 and 6 weeks, we observed histologic degeneration of the medial meniscus in all groups except the Sham group. In particular, the DMM and CATR groups showed a severe decline in the anterior horn. However, there was no significant difference in meniscus scores at 4 and 6 weeks.

**Figure 2.**
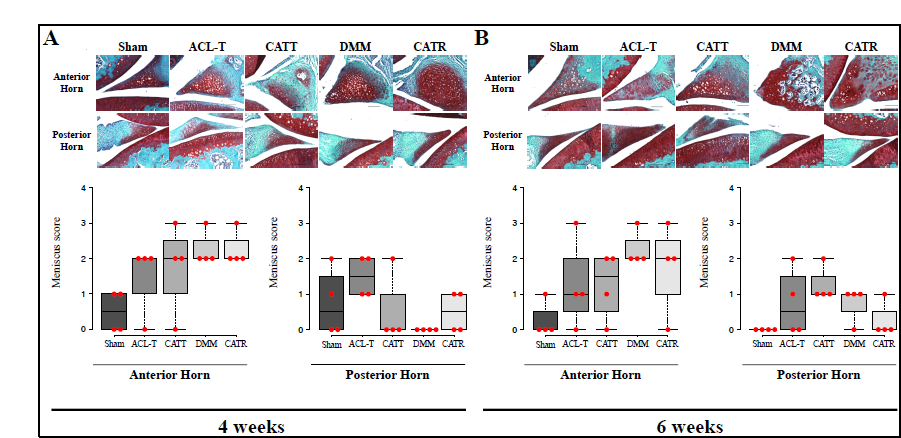
Histological analysis of the meniscus (A) Representative histology of the meniscus at 4 weeks and scoring results. Except for the Sham group, there were degenerative findings in the anterior horn of the meniscus, but there was no significant difference in the meniscus score. (B) Representative histology of the meniscus at 6 weeks and scoring results. Although the same degenerative findings were seen as at 4 weeks, there was no significant difference in meniscus score.

### μCT analysis of subchondral bone

To elucidate the characteristics of the subchondral bone within the model, we compared the subchondral bone changes within the model.

First, a representative image of the contact area in the sagittal and forehead planes for each model is shown in Figure 3. In the sagittal plane, the contact area of the ACL-T group was displaced posteriorly compared to the other groups. In the frontal plane, there was no characteristic difference in the contact area, but the medial meniscus of the DMM and CATR groups showed deviation (Figure 3, **white arrows**).

**Figure 3.**
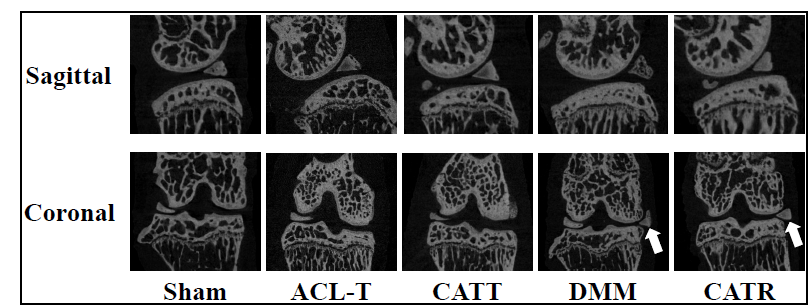
Reconstructed image of the knee joint Representative reconstructed images of the sagittal and coronal planes for each group. In the sagittal plane, the contact area of the ACL-T group was displaced posteriorly. In the coronal plane, we observed that the deviation of the medial meniscus in the DMM and CATR groups.

Next, we showed the results of the BV/TV in figure 4. There was no significant difference in the Sham group at 4 weeks and significantly reduced in the LA region compared with the medial posterior and posterior-lateral regions at 6 weeks (LA vs. MP, P =.010; LA vs. LP, P = .009). In the ACLT group, BV/TV was significantly increased in the MP region compared with the MA and LA regions at 4 weeks (MP vs. MA, P =.002; MP vs. LA, P < .001), and significantly increased in the MP region compared with the LA region at 6 weeks (MP vs. LA, P =.036). In the CATT group, at 4 weeks, BV/TV was significantly increased in the MP region compared with the MA and LA regions (MP vs. MA, P =.008; MP vs. LA, P = .001) and significantly decreased in the LA region compared with the LP region (LA vs. LP, P =.018). At 6 weeks, BV/TV was increased significantly in the MP region compared with the MA and LA regions (MP vs. MA, P =.045; MP vs. LA, P = .003). In the DMM group, BV/TV was significantly increased in the MP region compared with other regions at 4 weeks (MP vs. MA, P =.009; MP vs. LA, P < .001; MP vs. LP, P = .001). At 6 weeks, BV/TV was significantly increased in the MP region compared with other areas and significantly decreased in the LA region compared with the LP region (MP vs. MA, P <.001; MP vs. LA, P < .001; MP vs. LP, P < .001; LA vs. LP, P = .005). In the CATR group, at 4 weeks, BV/TV was significantly increased in the MP region compared with the LA and LP regions and significantly increased in the MA region compared with the LA region (MP vs. LA, P <.001; MP vs. LP, P = .014; MA vs. LA, P = .018). At 6 weeks, BV/TV was significantly decreased in the MP region compared with the LA region (MP vs. LA, P =.001).

**Figure 4.**
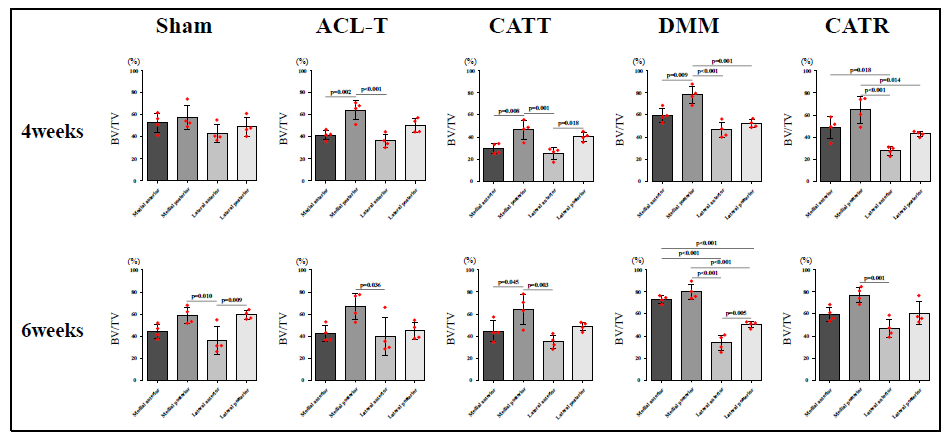
Analysis of subchondral bone within the model Results of BV/TV comparison within a model. The Sham group had a low LA area at 6 weeks. The ACL-T and CATT groups showed higher values in the MP region. The DMM and CATR groups tended to show higher MA and MP regions.

Subsequently, we performed comparisons between models for each region. We showed the results of the BV/TV in figure 5. In the MA region, at 4 weeks, the ACL-T and CATT groups showed a decreasing trend, with the ACL-T group significantly lower than the DMM group, and the CATT group significantly lower than the Sham, DMM, and CATR groups (ACL-T vs. DMM, P = .020; CATT vs. Sham, P = .004; CATT vs. DMM, P < .001; CATT vs. CATR, P = .017). At 6 weeks, The DMM and CATR groups were significantly higher than the Sham and ACL-T groups, and the DMM group was significantly higher than the CATT group (DMM vs. Sham, P < .001; DMM vs. ACL-T, P < .001; CATR vs. Sham, P = .047; CATR vs. ACL-T, P = .022; DMM vs. CATT, P < .001). In the MP region, at 4 weeks, the DMM group was significantly higher than the CATT group (DMM vs. Sham, P = .003). At 6 weeks, there was no significant difference. In the LA region, at 4 weeks, the CATT group was significantly lower than the Sham and DMM groups, and the CATR group was significantly lower than the Sham and DMM groups (CATT vs. Sham, P = .010; CATT vs. DMM, P = .002; CATR vs. Sham, P = .026; CATR vs. DMM, P = .006). At 6 weeks, there was no significant difference. In the LP region, at 4 weeks, there was no significant difference. At 6 weeks, the CATR group was significantly lower than the ACL-T group (CATT vs. Sham, P = .022).

**Figure 5.**
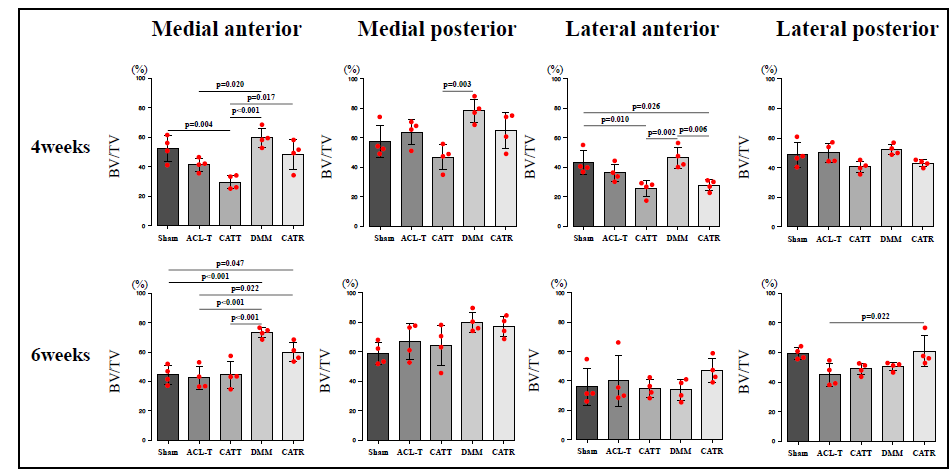
Analysis of subchondral bone between the model Results of BV/TV comparison between models. In the MA region, the CATT group decreased at 4 weeks and the DMM and CATR groups increased at 6 weeks.

## DISCUSSION

The different joint instability models showed various subchondral bone changes.^13^ However, differences in the region-specific characteristics of the subchondral bone remain unclear. In this study, we identified differences in the characteristics of meniscus degeneration and regional subchondral bone changes in different PTOA models. Meniscus degeneration was a trend observed in the OA model, but no significant differences existed between models. The region-specific subchondral bone changes in each model showed that changes in mechanical stress due to ACL and meniscus dysfunction, as well as changes in the contact area, affect the bony structure of the subchondral bone differently in each region (Fig. 6). Our results suggest that the changes in subchondral bone differ depending on the increase or decrease in mechanical stress at each region.

**Figure 6.**
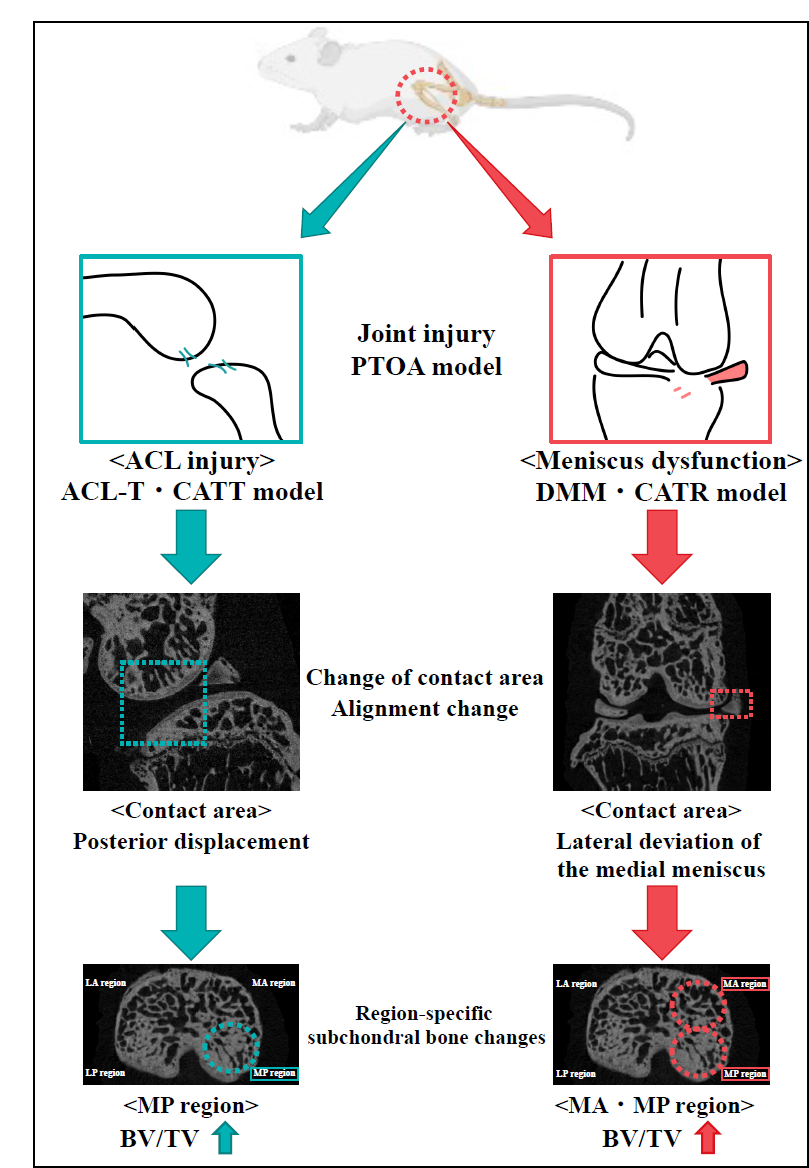
Summary of the results of this study The PTOA model causes different contact area changes and causes region-specific characteristics of the subchondral bone changes.

Previous studies have shown that meniscus lesions are a risk factor for OA and that a degenerated meniscus contributes to the progression of OA.^17–20^ In addition, it is clear that meniscus degeneration contributes to cartilage degeneration, but its effect on subchondral bone is unclear. Our histological analysis confirmed meniscus degeneration in all models except the Sham group, but there was no significant difference in scores. In our previous study, we observed meniscus degeneration in DMM and CATR at 8 weeks, but not significantly different from that in the Sham group.^16^ Thus, it is possible that the results of this study did not detect differences in meniscus degeneration between models because of the early time point. Furthermore, these results suggest that the differences in subchondral bone changes in the present study are not due to meniscal degeneration but increased mechanical stress due to kinematic and contact area changes associated with ACL and meniscal dysfunction.

Previous studies have shown that articular cartilage and subchondral bone properties vary from region to region.^14, 21–23^ In this study, we performed micro-CT analysis to elucidate the differences in subchondral bone changes in the different models. Evaluation of the contact area revealed a shift of the contact area posteriorly in the ACL-T group, suggesting that ACL functions inhibit anterior displacement of the tibia. Therefore, ACL transection caused the anterior displacement of the tibia, resulting in a change in the contact area. In addition, deviation of the medial meniscus was observed in the DMM and CATR groups. This is consistent with our previous study.^13^ The meniscus provides structural stability and dispersion of compressive stress to the knee joint. Therefore, it is speculated that increased compressive stress and instability due to semilunar deviation occur in the medial region of the DMM and CATR groups.

Articular cartilage has different mechanical properties depending on the region^14^, and mechanical stress induces cartilage degeneration. Therefore, subchondral bone may also have different mechanical properties at different regions depending on their mechanical environment. This study revealed differences in the characteristics of each region of the subchondral bone in different models. In the Sham group, BV/TV in the LA region was lower than in the MP and LP regions at 6 weeks. This trend was also observed in the other models. All models used in this study created bone tunnels in the femur and tibia. Therefore, the effect of the creation of bone tunnels may be canceled. In the ACL-T group, BV/TV in the MP region was higher than that in the MA and LA regions at 4 weeks, and at 6 weeks, BV/TV in the MP region tended to be higher than that in the other regions. Previous studies have shown that cartilage degeneration in the ACL-T model occurs in the posterior-medial region of the tibia.^24^ In addition, it is clear that the ACL-T model results in an anterior tibial translation.^13, 25, 26^ The results of this study also showed a posterior displacement of the contact area in the ACL-T group (Fig3). Therefore, it is suggested that the increase in mechanical stress associated with the change in contact area after ACL transection caused the increase in BV/TV in the MP region. In the CATT group, as in the ACL-T group, BV/TV in the MP region was higher than that in the MA and LA regions at 4 weeks, and at 6 weeks, BV/TV in the MP region was still higher than that in the MA and LA regions. The CATT model suppresses abnormal tibial translation of the ACL-T model. However, previous reports have shown that the CATT model has increased abnormal tibial translation compared to the Sham model.^13^ Therefore, the subchondral bone changes observed in the CATT group may be related to changes in mechanical stress due to displacement of the contact surface. In the DMM group, BV/TV in the MP region at 4 weeks was higher than that in the other regions. At 6 weeks, BV/TV tended to be higher in the MP and MA regions. The DMM model induces OA due to meniscus dysfunction. Previous studies have reported that increased compressive stress and joint instability may be responsible for cartilage degeneration in the DMM model.^27–29^ Furthermore, previous studies have reported that BV/TV in the DMM model increases in the early stages.^30^ In the results of this study, deviation of the medial meniscus was observed in the DMM group, and the subchondral bone changes in the medial region may be caused by the increased compressive stress associated with meniscus deviation. In the CATR group, the MA and MP regions showed higher BV/TV at 4 weeks than the LA region, and the MP region had higher BV/TV at 4 weeks than the LP region. The CATR model is a model that suppresses the joint instability of the DMM model. Previous studies have confirmed lateral deviation of the medial meniscus and meniscus degeneration in the CATR model as well as in the DMM model.^13, 16^ Therefore, an increase in compressive stress is inferred in the CATR model as well as in the DMM model, which may have caused changes in the subchondral bone in the medial region.

We have previously reported differences in subchondral bone changes between different mechanical stress models.^13^ However, the differences in the characteristics of each region still need to be clarified. In this study, we compared the characteristics of different regions between models. In the MA region, at 4 weeks, BV/TV in the CATT group decreased compared with the other groups except for the ACL-T group, and at 6w weeks, BV/TV in the DMM and CATR groups increased compared with the other groups. The decrease in BV/TV in the CATT group is consistent with our previous reports.^13^ However, the region-by-region analysis of this study suggests that changes in the contact areas may be responsible for the decrease in BV/TV in the MA region in the ACL-T and CATT groups. The DMM and CATR groups are both models of meniscus dysfunction, and previous studies have confirmed that meniscus dysfunction increases compressive stress.^28, 29^ In addition, deviation of the meniscus was also observed in both models (Fig3). Thus, it is suggested that an increase in compressive stress may increase BV/TV. These results suggest that ACL deficiency and meniscus dysfunction, which cause PTOA, have different effects on subchondral bone changes. In the lateral region, BV/TV in the CATT and CATR groups in the LA region was reduced compared to the Sham group, and a similar trend was observed in the LP region. The CATT and CATR models suppressed joint instability. Therefore, surgical intervention to suppress joint instability may have affected the results. Previous studies have confirmed that the CATT and CATR models do not limit a range of motion.^13^ However, the CATT and CATR models may limit a dynamic range of motion, which needs to be examined in the future.

There are two significant limitations to this study. In this study, we analyzed at 4 and 6 weeks after surgical intervention. The reason for this was to reveal changes in the meniscus and subchondral bone in the early stages of OA. Therefore, we have not been able to evaluate changes over the later stages of OA. Future studies will be required to validate additional longer time points. The effects of bone tunneling and suppression of joint instability on knee joint motion in mice are unclear. We have confirmed that suppression of joint instability and creation of bone tunnels do not cause limitations in joint range of motion. However, we have not been able to evaluate them during cage activities such as walking. Future studies should additionally examine this point and clarify the effects of the suppression of joint instability on joint motion.

## Conclusions

In conclusion, this study showed that subchondral bone changes in different models of mechanical stress were different in each region. In particular, changes in the contact area and increased compressive stress due to meniscus dysfunction were suggested to promote bone formation. It has been shown that a decrease in articular cartilage thickness is associated with alignment changes in knee OA patients.^31^ The results of this study indicate that changes in alignment and contact area in the PTOA model may cause region-specific characteristics of the subchondral bone changes. Further studies will be required to elucidate the relationship between articular cartilage degeneration and subchondral bone changes caused by alignment changes, and to identify early-stage knee OA lesions and prevent the progression of knee OA.

## Author Contributions

All authors approved the final version to be published.

Study design: KA, YU, and TK

Data collection, Histological analysis: KA and KT

Subchondral bone analysis: KA and KT

Manuscript composition: KA and TK

## ACKNOWLEDGEMENTS

The author(s) received no financial support for the research, authorship, and/or publication of this article.

## DECLARATION OF CONFLICTING INTERESTS

The authors declare that there is no conflict of interest

### Ethical Approval

This study was approved by the Animal Research Committee of Saitama Prefectural University (Approval Number: 2020-1).

### Animal Welfare

This study followed institutional guidelines for humane animal treatment and complied with relevant legislation.

